# Inferring quantity and qualities of superimposed reaction rates in single molecule survival time distributions

**DOI:** 10.1101/679258

**Authors:** Matthias Reisser, Johannes Hettich, Timo Kuhn, J. Christof M. Gebhardt

## Abstract

Actions of molecular species, for example binding of transcription factors to chromatin, are intrinsically stochastic and may comprise several mutually exclusive pathways. Inverse Laplace transformation in principle resolves the rate constants and frequencies of superimposed reaction processes, however current approaches are challenged by single molecule fluorescence time series prone to photobleaching. Here, we present a genuine rate identification method (GRID) that infers the quantity, rates and frequencies of dissociation processes from single molecule fluorescence survival time distributions using a dense grid of possible decay rates. In particular, GRID is able to resolve broad clusters of rate constants not accessible to common models of one to three exponential decay rates. We validate GRID by simulations and apply it to the problem of in-vivo TF-DNA dissociation, which recently gained interest due to novel single molecule imaging technologies. We consider dissociation of the transcription factor CDX2 from chromatin. GRID resolves distinct, decay rates and identifies residence time classes overlooked by other methods. We confirm that such sparsely distributed decay rates are compatible with common models of TF sliding on DNA.

## Introduction

The actions of biomolecules are governed by thermal fluctuations and thus are intrinsically stochastic. Accordingly, interactions such as association and dissociation events of molecular species often follow Poissonian statistics with a constant probability per time, the rate constant, to occur, and the experimentally accessible reaction lifetimes are exponentially distributed. Generally, not all members of a molecular species undergo the same type of interaction at any time, but each conducts one of multiple possible types of interaction. If a measurement to determine reaction lifetimes cannot distinguish between these different reaction types, the resulting lifetime distribution is a Laplace transform of a spectrum of reaction rates from superimposed Poisson processes, and thus multi-exponential (Figure 1a). Retrieving the underlying reaction rate spectrum consequently evokes an inverse Laplace transformation.

**Figure 1:**
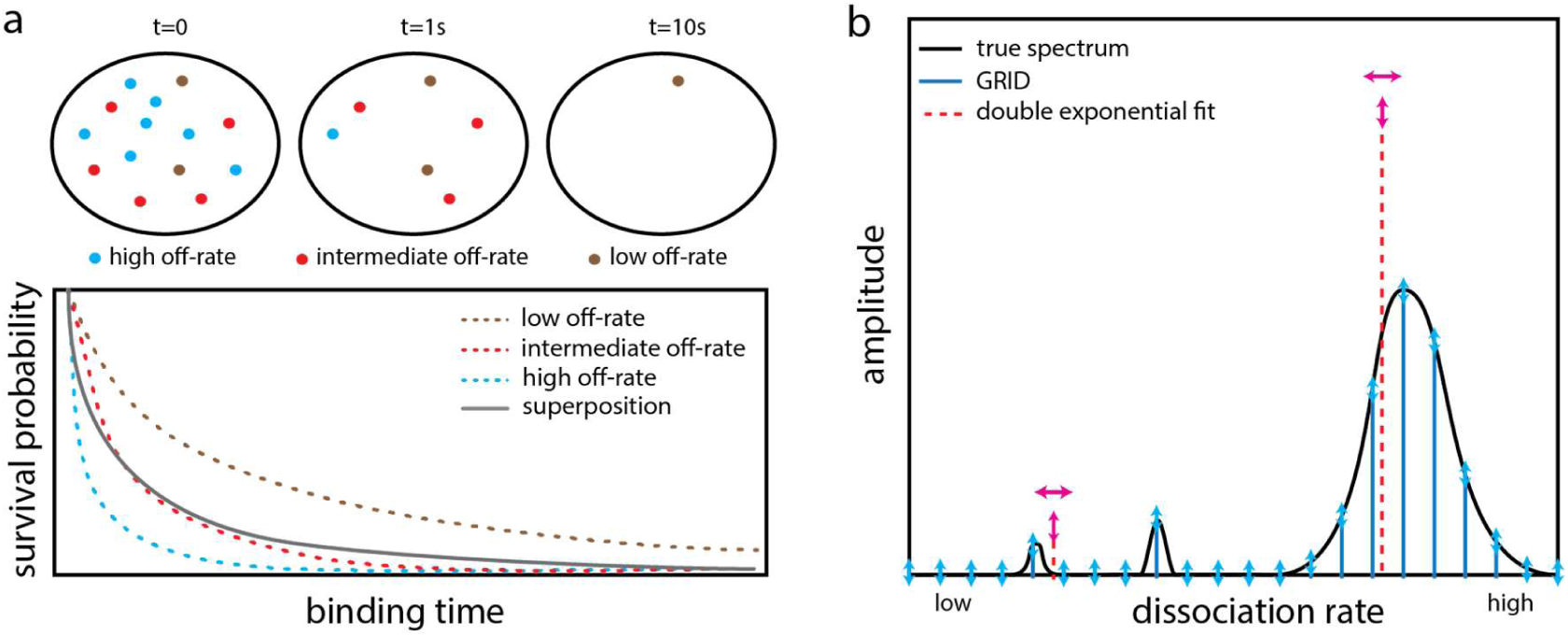
Working principle of GRID. (a) Sketch of a TF exhibiting three distinct dissociation processes from chromatin (upper panel). The resulting survival time distribution is a superposition of the three processes (lower panel). (b) Sketch of a decay rate spectrum (black solid line) underlying a complex survival time distribution. In common multi-exponential analysis the decay rates and their amplitudes are varied (red dashed lines). In contrast, GRID only varies the amplitudes of a grid of decay rates (blue solid lines). Degrees of freedom are indicated by arrows.

The inverse Laplace transformation is an ill-posed problem for inherently noisy, discrete distributions and numerical solutions are often unstable ^1, 2^. Nevertheless, algorithms treating the Laplace transform using gradient methods and appropriate regularization have been successfully developed for noisy data in NMR ^3, 4^ and protein folding ^5^. An elegant method based on phase functions avoids fitting procedures and enables direct reconstruction of the rate spectrum of superimposed ^6^ and sequential ^7^ biological decay processes.

Lifetimes of biomolecular interactions are frequently measured by single-molecule fluorescence microscopy ^8–16^. In such experiments, photobleaching of the fluorescent label adds a parallel decay path to each reaction type but is indistinguishable from a successful reaction ^17^. This complex kinetic scenario cannot be solved by the phase function method or current approaches of numerical inversion of the Laplace transform. Survival time distributions are for example corrected for by comparison to immobile molecules such as histone H2B ^11, 18^ or the photobleaching rate constant is directly considered using several time-lapse conditions ^9, 19^.

An example for superimposed reactions are transcription factor (TF) – chromatin interactions. TFs may be involved in a manifold of different binding reactions, such as binding to specific or numerous different unspecific sequences on either free or nucleosomal DNA, binding to RNA or to other proteins. To circumvent Laplace inversion methods, current analysis of TF – chromatin interactions apply models with a fixed number of exponential functions to describe distributions of fluorescence survival times by varying the decay rates and their amplitudes ^8, 10, 11, 20^. Such exponential fitting is robust but requires knowledge of the number of decay rates and thus is limited when resolving complex decay rate spectra.

Here, we tackle the problem of inverting the Laplace transform in the context of fluorescence survival time distributions obtained by single molecule imaging subject to photobleaching. We apply a grid of invariable decay rates and fixed spacing with variable positive weights to describe distributions of fluorescence survival times. Reducing the number of nonlinear parameters and specializing a regularization enables us to robustly use gradient methods to infer rate spectra. We validate our genuine rate identification (GRID) method by simulations and show that GRID enables inferring complex rate spectra of superimposed interactions with both narrow and wide distributions of decay rates. We apply GRID to analyse fluorescence survival time distributions of dissociation events between the transcription factor CDX2 and chromatin recorded in live cells. GRID extends the information obtained by multi-exponential fitting approaches. In particular, in contrast to intuitive expectation with numerous different bound DNA sequences in mind, we do not observe a broad cluster of unspecific dissociation rates. A distinct unspecific decay rate is consistent with a quantitative solution to a common model of TFs sliding on and binding to unspecific DNA sequences.

## Results

### Analysing superimposed reactions by GRID

We considered several parallel reactions each following Poissonian statistics with distinct dissociation rates giving rise to exponentially distributed lifetimes in the time domain (Figure 1a). We further considered measurements of reaction lifetimes by single molecule fluorescence microscopy using fluorescent labels subject to photobleaching. The corresponding survival time distribution is a superposition of exponential functions weighted by the relative occurence of the respective process and enveloped by the decay of fluorescent labels (Figure 1a, Methods, Equation 2).

Solving the inverse Laplace transformation to infer the number and amplitudes of reaction processes from this multi-exponential distribution is impeded by the presence of the photobleaching process. We reduced the number of non-linear parameters in a corresponding minimization problem by introducing a grid of densely spaced invariable decay rates (Figure 1b and Methods). We further designed a cost function restricting the amplitudes to positive values for physical reasons. The cost function also accounted for the integration time of the camera, which limits the time resolution of fast dissociation rates. We refer to this regularization as mean decay regularization (MDR) (Methods). If this regularization is omitted, fast decay rates are used to account for noise in the first data points of the time-lapse records without compromising overall fit quality, since fast dissociation rates introduce negligible error at large times. The cost function can accommodate any number of time-lapse conditions in single molecule fluorescence measurements. We used the gradient method implemented in the Matlab fmincon function to solve the minimization problem corresponding to the inverse Laplace transform (Methods).

We validated our approach, GRID, using simulated survival time distributions. We simulated distributions as would be obtained by single molecule fluorescence measurements with up to 10 time-lapse conditions spanning a range between 0.05 s and 50 s and 10.000 recorded reaction events per condition (if not stated otherwise), a photobleaching rate constant *k* = *1* s^−1^ and considered noise intrinsic to Poisson processes (Methods and Supplementary Table). To test the performance of GRID, we compared different cost functions and varied several qualities of the rate spectra including the quantity of well-separated superimposed rates, rate values and amplitudes and the width of densely spaced rate clusters.

First, we compared the performance of the MDR regularization in our cost function to generic regularizations such as the L2-norm ^21^ and the L4-norm of the fitted parameters and a more specific norm that weights fitting parameters with the hyperbolic cosine (Methods). We simulated survival time distributions (1000 events per time-lapse condition) with two superimposed reactions with rates of 0.1 s^−1^ and 5 s^−1^ (Figure 2a). While all alternative regularizations showed artificial broadening of rate distributions, our MDR regularization successfully reproduced the ground truth rate spectrum (Figure 2a and b). We thus retained our cost function for the remainder of the study.

**Figure 2:**
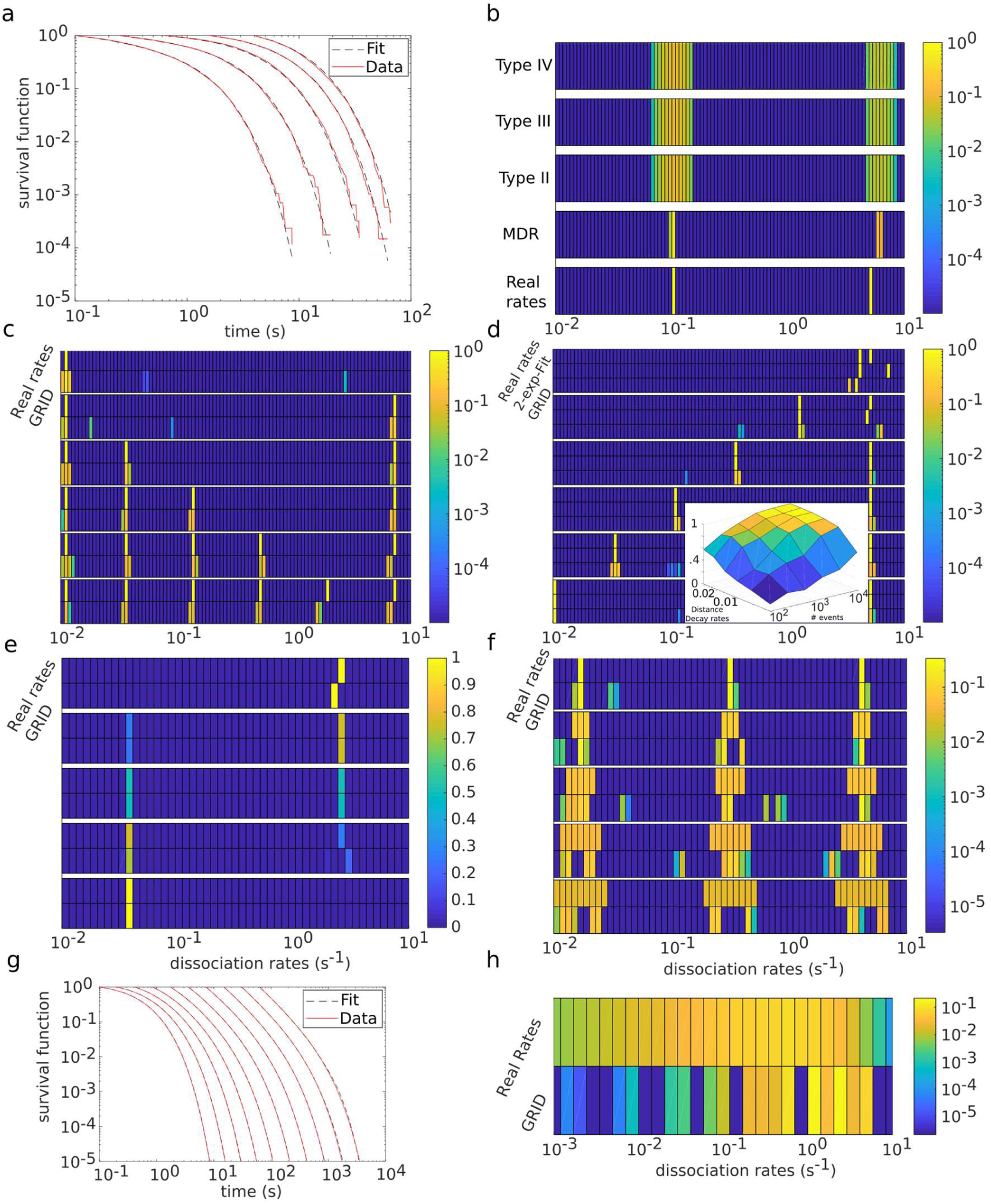
Validation of GRID. (a) Simulated survival time distributions with dissociation rates of 0.1s^−1^ and 5s^−1^ and photobleaching rate of 1s^−1^ (red lines) and distributions obtained using the results by GRID displayed in b (black dashed lines). (b-f,g) Heat maps comparing the ground truth rate spectrum used to simulate survival time distributions and the rate spectrum obtained by GRID. Simulations include a photobleaching rate of 1s^−1^. Amplitudes are color coded. (b) Comparison of different regularizations (specified in Methods) used in the cost function of GRID. (c) Increasing number of distinct decay rates starting at k = 0.011s^−1^ and separated by a factor of 4. (d) Increasing separation between two distinct decay rates. *k*_fast_ = 5 s^−1^ and *k*_slow_ in [0.01,4] s^−1^. Inset: influence of the number of detected events and separation of decay rates on the accuracy of the inferred spectrum. (e) Variable amplitudes of two distinct decay rates (*k*_slow_ = 0.035 s^−1^ and *k*_fast_ = 2.44 s^−1^). (f) Increasing width of three decay rate clusters centred at *k*_slow_ = 0.016 s^−1^, *k*_int_ = 0.3 s^−1^ and *k*_fast_ = 3.9 s^−1^. Relative width is up to 70%. (g) Simulated survival time distributions of a power-law (specified in Methods) and photobleaching rate of 1s^−1^ (red lines) and distributions obtained using the results by GRID displayed in h (black dashed lines). (h) Power-law distribution (specified in Methods).

Second, we simulated survival time distributions with an increasing number of superimposed reactions with rates between 0.01 s^−1^ and 10 s^−1^, separated by at least a factor of 4. Within this range and spacing, GRID reliably identified up to six distinct reaction rates (Figure 2c). False positive rate detections only appeared as minor background in the spectra.

Third, we investigated whether the spacing of rates influenced the performance of GRID. We simulated survival time distributions with a fast dissociation rate fixed at 5 s^−1^ and varied a slow dissociation rate between 10^−2^ s^−1^ and 4 s^−1^ (Figure 2d). GRID inferred rate values reliably up to a separation by a factor of ∼2, consistent with the resolution limit of exponential analysis ^22^ and comparable to a two-exponential fit. Analogously, we varied the fast dissociation rate between 10^−2^ s^−1^ and 10 s^−1^ while keeping the slow dissociation rate constant at 5.4·10^−3^ s^−1^ (Supplementary Figure 1). Again, the values of both rates were accurately determined up to a separation by a factor of ∼2, and comparable to a fit by a double-exponential model.

To estimate the influence of the number of simulated reaction events on the accuracy of the inferred rate spectra for the case of two simulated dissociation rates with variable spacing, we varied the number of simulated reaction events between 100 and 10.000 per time-lapse condition and quantified the deviation from the ground truth using 100 independent simulations (Figure 2d and Methods). In line with ^23^, the closer the reaction rates, the more reaction events need to be observed to resolve them.

Fourth, we examined the response of GRID to the amplitudes of reaction rates. We simulated survival time distributions with two rates of 0.035 s^−1^ and 2.44 s^− 1^ and varied their amplitudes from 0% to 100% (Figure 2e). Since rates identified by GRID oftentimes split between two grid positions including and next to the ground truth (Figures 2b-d), we binned grid positions for the analysis of amplitudes (Methods). GRID well recovered both rates and their amplitudes.

Fifth, we tested to which extend GRID was able to resolve rate spectra of more complex shape. Thus, we simulated survival time distributions (100.000 events per time-lapse condition) using three dense square shaped decay rate clusters at centre positions of 0.016 s^−1^, 0.3 s^−1^ and 3.9 s^−1^ and stepwise increased their width from 0% to 70% relative width (Figure 2f). GRID recovered the width of rate clusters in most scenarios. However, a tendency to split clusters into two close sub clusters became apparent.

Since a power-law behaviour of TF – chromatin dissociation has been suggested ^14, 24^, we tested whether GRID would accurately resolve a power-law shaped ground truth. In principle, GRID is able to handle power-law distributions (Methods). We simulated a survival time distribution (100.000 events per time-lapse condition) corresponding to a power-law including photobleaching and noise (Methods, Equation (10)) (Figure 2g). GRID split the broad distribution of decay rates into sub clusters (Figure 2h). However, the resulting decay rate spectrum is well distinguishable from a sparse distribution of decay rates (Figure 2b-e).

### GRID analysis of CDX2 dissociation from chromatin

A current biological question of high interest is the interaction of transcription factors with chromatin. In particular, it is unclear how many kinetically distinct interactions a TF undergoes with chromatin. Thus, after having validated our rate analysis approach with simulations, we applied GRID to survival time distributions of CDX2 dissociation from chromatin, obtained by live-cell single-molecule tracking of the fusion protein Halo-CDX2 ^25^ labelled with SiR-dye (Methods and Supplementary Movie 1). We recorded survival time distributions of four time-lapse conditions (Figure 3a). GRID inferred a rate spectrum with five clearly distinct rates ranging from 5 s^−1^ to 0.006 s^−1^ with strongly decreasing amplitude and a photobleaching rate of 0.1 s^−1^ (Figure 3b). We did not observe any broad rate clusters. Simulated distributions using the dissociation rate spectrum extracted from the data by GRID well overlapped with the measured survival time distributions (Figure 3a), in contrast to survival time functions calculated using dissociation rates obtained by fitting a triple-exponential model (Figure 3a). Thus, compared to common multi-exponential analysis, dissociation rates inferred by GRID better describe the measurement.

**Figure 3:**
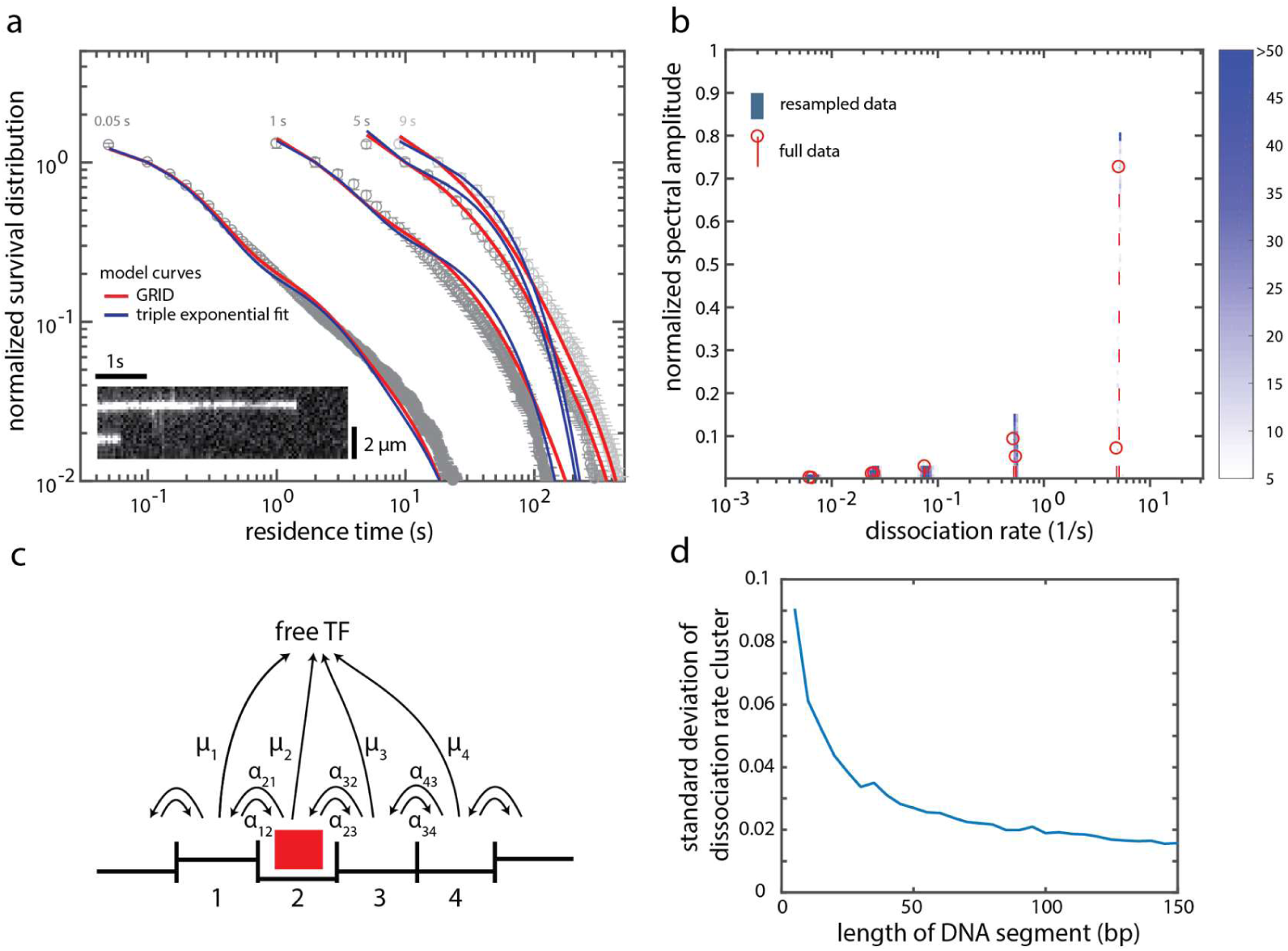
Dissociation rate spectrum of TF – chromatin interactions. (a) Fluorescence survival time distributions of SiR-Halo-CDX2 obtained by live-cell single molecule imaging (grey symbols), fit of tri-exponential model (blue lines) and distributions obtained using the results by GRID displayed in b (red lines). Time-lapse conditions are indicated above the distributions. Inset: Kymograph of two molecules detected within a 0.05s time-lapse movie. The graph contains data from 10653 molecules in 79 cells. Error bar denotes s.d. (b) The dissociation rate spectrum obtained by GRID using all data (red dashed lines and circles) and heat map of GRID results obtained by 499 times resampling 80% of data (blue color code) (c) State diagram of a TF (red box) diffusing on and dissociating from DNA. Each binding position is associated with an individual binding energy. The parameters are specified in Methods. (d) Width of decay rate cluster of unspecific TF – chromatin interactions obtained by solving the state diagram in c for 500 DNA segments as function of segment length (for details see Methods).

To test the extracted rate spectrum for consistency, we omitted the fastest time-lapse condition of 0.05s in the analysis, which exclusively contains temporal information between 0.05 and 1 s (Supplementary Figure 2a). As expected, the extracted rate spectrum is devoid of the dissociation rate at 5 s^−1^, while the remaining spectrum does not change significantly (photobleaching rate was 0.4s^−1^) (Supplementary Figure 2 b). When omitting the longest time-lapse condition of 9s, which contains similar temporal information to the time-lapse condition of 5s, the extracted rate spectrum does not change significantly, as expected (photobleaching rate was 0.1s^−1^) (Supplementary Figure 2c and 2d).

It is commonly assumed that dissociation occurs from a few specific sequences and a plethora of unspecific sequences including one or several mismatches at various positions. Intuitively, for dissociation from unspecific sequences, such a picture results in dissociation rates spreading over a broad range of values, potentially giving rise to a power-law distribution of decay rates ^14, 24^. To substantiate and challenge this intuitive picture of unspecific dissociation, we estimated the cluster width arising from unspecific TF – DNA dissociation using a well-accepted model of TF sliding on unspecific DNA with dissociation from any site within the sliding segment ^26–28^(Figure 3c and Methods). We found that the dissociation rate from a single segment would reduce to a single value due to fast 1D diffusion. When considering several segments, the corresponding dissociation rates combined to a narrow cluster, due to stochasticity in the base pair content of different segments. The width of the resulting decay rate cluster was anti-correlated with the length of the segments (Figure 3d). However, even unreasonably small sliding segments resulted in cluster widths well below the resolution of GRID. Overall, our calculations suggest that unspecific TF – DNA interactions eventually result in a single resolvable dissociation rate, consistent with our measurement.

## Discussion

### GRID reveals rate spectra underlying complex survival time distributions

We introduced GRID, an approach to extract reaction rates from experimentally measured fluorescence survival time distributions of complex superimposed reactions. GRID robustly identifies the number and amplitudes and gives information on cluster width of reaction rates, even if lifetime measurements are aggravated by photobleaching of fluorescent labels. Such distorting additional decay rates hamper the use of previously reported approaches to tackle the inverse Laplace transformation of survival time distributions (Barone et al., 2001, Berman et al., 2013, Voelz and Pande, 2012, Zhou and Zhuang, 2006).

GRID has the advantage that the number of decay rates in the biological system does not have to be guessed. This is a major drawback of current analysis schemes using a small number of decay rates. Our simulations suggest that GRID, despite being the more complex approach, does not come with a loss in accuracy in a situation where the number of decay rates is known. A second advantage of GRID is that it can reveal broad clusters of reaction rates, information intrinsically inaccessible to analysis schemes using a small number of decay rates. While we validated and applied GRID to data sets including photobleaching and several time-lapse conditions, it should in principle be applicable to individual survival time distributions already corrected for photobleaching.

GRID is currently restricted to superimposed reactions following Poissonian statistics with positive amplitudes. Thus, GRID is not applicable to arbitrary survival time distributions (Methods). Due to computation costs, the number of rates in the grid is currently limited to 200. Consequently, the resolution to identify decay rates is limited, and oftentimes GRID splits a single decay rate onto two adjacent grid positions. Compared to (Zhou and Zhuang 2006), GRID converts the inverse Laplace transformation into an optimization problem, with the accompanying disadvantage of a large number of degrees of freedom. This requires introducing a robust regularization. Additionally, a large number of measurements are advisable. While GRID allows differing distinct decay rates from broad clusters of decay rates or a power-law distribution, it is limited in identifying the shape of such clusters or distributions with high accuracy.

### Rates of CDX2 – chromatin dissociation

GRID resolves five distinct dissociation rates corresponding to chromatin residence times between 0.2 s and 170 s from the fluorescence survival time distributions measured for the dissociation of CDX2 from chromatin. Their amplitudes differ significantly, between ca. 80 % and < 5 %, pointing to a low occurrence of the slowest dissociation processes. Yet, long interactions appear necessary for a full description of the measured survival time distributions, as multi-exponential fitting using three dissociation rates as reported previously for different TFs ^20, 29^ failed to fully recover the measured survival time distributions. The fastest rate of TF – chromatin interactions was previously identified as binding of the TF to unspecific DNA sequences ^14, 30, 31^. Analogously, CDX2 might exhibit transient unspecific and stable specific binding to chromatin. A detailed assignment of rates to specific reaction mechanisms of CDX2 is a task for future studies.

In the presence of unspecific LacI – chromatin interactions, a power law was found to well describe the survival time distribution, potentially representing a multitude of co-occurring different dissociation rates ^14, 24^. For CDX2, we thus expected a broad cluster of densely spaced dissociation rates in the fast rate regime. However, despite the capability of GRID to resolve such clusters, we observed a single well-defined rate. This observation is strengthened by our model of TF – DNA interactions combining sliding and dissociation, where we found that differences in dissociation rates between several sliding segments are well below the resolution limit of GRID.

In principle, dissociation rate spectra can be converted to binding energy landscapes. Such an approach has been demonstrated for membrane proteins ^32^. In the case of TF – chromatin interactions, this approach is challenging. Our measurement and modelling suggest that unspecific dissociation rates from unconnected DNA segments are not resolvable. For binding to a specific target site adjacent to unspecific sequences, observable dissociation rates only reflect effective rates, not the real dissociation rate from the specific target site ^33^. Furthermore, amplitudes of measured dissociation rates need to be converted to occupation frequencies. In addition, recent measurements suggest that some TFs may form local condensates ^34^. Thus, density of states, or energy states if the reaction coordinate is known, can only be calculated with additional knowledge of the underlying microscopic mechanisms. Finally, a comprehensive picture should also consider the relative distributions of diffusive and bound states, which can for example be obtained from comprehensive modelling kinetics ^35, 36^ or interlaced time lapse measurements ^37^.

## Materials and Methods

### Model for the survival time function of an ensemble of chromatin-bound fluorescently labelled TFs

We assume that dissociation of a TF from any bound state, in particular from a bound DNA sequence, follows Poissonian statistics with a dissociation rate constant *µ*_*i*_ characteristic for this particular state. We further assume that the TF may bind to a multitude of different DNA sequences, both unspecific and specific. The probability of a particular dissociation event to occur be *S*_i_.

For independent dissociation processes, the resulting survival time function of an ensemble of TFs is a superposition of individual dissociation processes

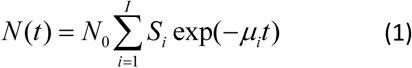

for the remaining bound population *N* at time t if *N*_*0*_ TFs were bound at time *t=0. N*_*0*_*·S*_*i*_ is the number of TFs in the ensemble that exhibit the dissociation rate constant *µ*_*i*_. The total number of dissociation processes is denoted by *I*.

So far, we assumed that the survival time function of bound TFs is only determined by dissociation. However, in single molecule fluorescence experiments, the TF is identified by a fluorescent label prone to photobleaching. Thus, the experimentally observed termination of a bound state may be due to photobleaching of the fluorescent label or dissociation of the TF. We assume that photobleaching again follows Poissonian statistics.

The fluorescence survival time function observed in experiments then reads

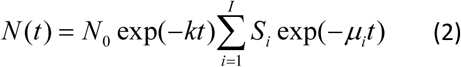

where *k* is the photobleaching rate constant. According to this equation, only the sum *k*+*μ*_i_ can be inferred from the fluorescence survival time distribution. To separate photobleaching from dissociation, we perform time-lapse measurements ^9^.

### Simulation of TF dissociation kinetics

We simulated survival time distributions of TFs with effective dissociation rate constants *k*_*eff,i*_ = a /*τ*_*tl*_ + *k*_*off,i*_ accounting for dissociation with dissociation rate constant *k*_*off,i*_ and with the photobleaching number a and the time-lapse period *τ*_*tl*_. Different *k*_*off,i*_ occurred with probability *S*_*i*_. We first generated a random number with uniform distribution to draw the *k*_*eff,i*_ from the probability distribution S. Next, we generated a new random number and transformed it to an exponential distribution with the constant *k*_*eff,i*_ to obtain the time at which the TF dissociated. This time entered a distribution with a bin-size corresponding to the time-lapse period. We repeated this procedure N times to obtain a survival time distribution of N TFs. To obtain a complete dataset, we repeated this procedure for various time-lapse periods *τ*_*tl*_. Simulations were conducted in MATLAB.

### A Method for the inverse Laplace transform

To determine the dissociation rate spectrum of the TF, we could in principle fit the fluorescence survival time function including bleaching, Equation 2, to the measured distributions obtained from several time-lapse conditions. To ensure robust inference of dissociation rates, we reduce the number of free parameters by applying a grid of I invariable dissociation rates with fixed spacing and numerically determine the probabilities Si of each dissociation rate. Summing up, the number of unknown parameters is *I*+1, namely [k, S]. Since *I* is usually larger than the number of observables in time-lapse measurements, the fitting problem is underdetermined. Thus, to obtain a unique solution, we apply regularizations based on basic physical considerations and time resolution constraints of the measurement process.

As first regularization, we introduce the constraint *S* ≥ 0 of non-negative probabilities and *k* ≥ 0 of a non-negative photobleaching rate constant. This ensures our model is monotonically decreasing, as expected from superimposed Poisson processes. As second regularization, we account for the integration time τ_int_ of the camera used to record fluorescent light. This integration time limits the time resolution of fast dissociation rates (*µ_i_* > *τ*_*int*_ ^−1^). As a mathematical measure for this limitation, we introduce the time dependent expectation value of the dissociation rate <*µ*> of the bound TF population

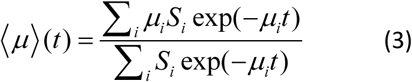

where the value 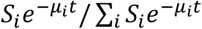 may be interpreted as the time dependent probability to find a TF that exhibits the dissociation rate µi at time t. We then introduce the expression

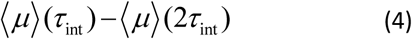

to describe the change of the mean dissociation rate in the dead time of our measurement. By minimizing this quantity, we reduce the number of degrees of freedom during our dead time and thereby avoid overfitting. We refer to this regularization as the mean decay regularization (MDR).

We next define the difference between the fluorescence survival time function and the measured distribution of the m-th point in the n-th time-lapse record, *Δf*_*nm*_, as

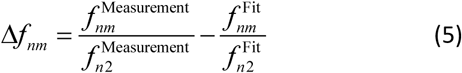

where we normalized the values of the fitted and measured distributions to the population at the second time point of a time-lapse record to eliminate the unknown amount of the initial population.

We further introduce the cost function L of the fitting problem which consists of the difference between measurement and theoretical function, *Δf*_*nm*_, and the regularization of the mean dissociation rate

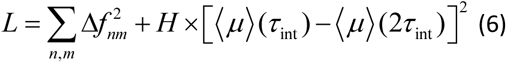

Since both *Δf*_*nm*_ and the regularization contribute to the same cost function, we introduce the empirical parameter H to limit the influence of the regularization.

The complete optimization problem finally is

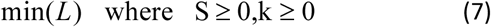

We solve this optimization problem with the gradient based method fmincon solver with the sequential quadratic programming algorithm of the Matlab optimization toolbox, to find the spectrum of dissociation rates of TF-chromatin dissociation. We refer to our cost function as MDR.

Alternatively, we tested the cost functions of the L2-norm (Type II) 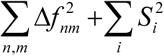, the L4-norm (Type III) 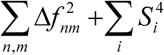and a more specific norm that weights fitting parameters with the hyperbolic cosine 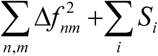 cosh(0.1·*k_i_*) (Type IV).

### Application of GRID to power-law functions

In GRID, we restricted ourselves to positive off-rates, positive amplitudes and a positive bleaching rate. Therefore, GRID can be applied to a certain type of model-functions. The model functions as well as the absolute value of their derivatives has to decay strictly monotonously. In particular we show here that GRID can be applied to power-law models.

We construct a survival function by calculating the power-law

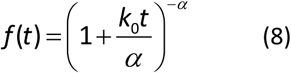

where *k*_0_ is a constant that shifts the pole to *t* < 0. The number *α* needs to be larger than one so that the average binding time of the TF converges. This model converges to a single exponential function in the limit *α* →∞. We analytically calculated the spectrum *S(k)* of (8) as

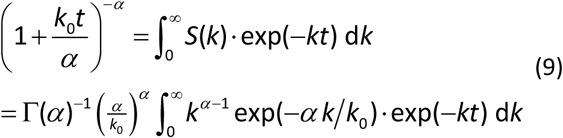

To check if the time-lapse approach combined with GRID can recover this power-law we calculated a survival time distribution according to

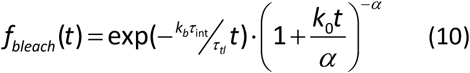

To introduce noise we stochastically resampled this survival function.

### Model for observable dissociation rates

In our model of TF-DNA dissociation, we assume that the TF binds to a free segment of DNA with a length corresponding to N base pairs. Within the DNA segment, the TF can assume N different binding positions (Figure 3, Inset). The TF slides between binding positions within this segment by 1D diffusion, restricted by roadblocks at the edges of the segment ^38, 39^. The TF can leave the segment by dissociating from any position within the DNA segment. We further consider the variance σ of DNA binding energies in units of *k*_*b*_*T*. This energetic variance leads to dissociation rates that are normal distributed around the mean dissociation rate *μ* with a standard deviation *σ*. The variance of unspecific binding energies was previously estimated to be *σ* <= 1 *k*_*b*_*T* ^40^. To each TF position within the DNA segment, we ascribe a random dissociation rate from this distribution.

The rate of sliding of the TF from state (or position) i to j be α_ij_. The ratio of α_ij_ and α_ji_ is determined by the energy difference between the two positions, which in turn is determined by the dissociation rates of the TF from DNA at positions i and j. To find values for α_ij_ and α_ji_, we assume that the transition rate to a lower binding energy level is given by the diffusion rate, while the transition to a higher binding energy level is limited by the energetic gap between the two levels. With this assumption we calculate the transition rates according to the law of detailed balance

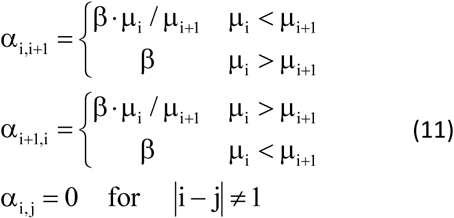

where β is a mean sliding rate. As described in ^33^ the Kolmogorov formalism can be used to model the dynamics of the TF on DNA. We find the time-dependent probability *p_i_* of the TF to be in state *i*

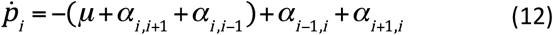

The observable dissociation rates are determined by the eigenvalues of the eigenvalue-problem and their amplitudes in the solution for the time dependent probability. We calculate these amplitudes by introducing the initial condition 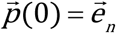, which is the unitary vector in n-th direction. This initial condition states the initial position of the TF after association to the DNA segment.

We assume a mean sliding rate of β= 10^+4^, calculated based on in vivo sliding measurements ^28, 33^. Since β is much larger than the fastest dissociation rate typically measured in experiments (< 10 s^−1^), the amplitudes of all except one eigenvalue become negligible for a single sliding segment.

To describe overall TF binding in the nucleus, we consider 500 independent unspecific DNA segments. As above, each segment contributes a single dissociation rate corresponding to the particular dissociation rate distribution of this segment of DNA. Due to stochastic fluctuations of dissociation rates drawn for each segment, the mean dissociation rates of different segments form a narrow cluster of dissociation rates.

### Quantitative comparison between rate spectra

To quantify the resemblance between inferred spectrum and ground truth in Figure 2d and Supplementary Figure 1, we calculated the scalar product of these two spectra. This value is high if the rates are at the same position and low if the rates are shifted with respect to each other. The scalar product is zero if the rates are shifted by one increment in the GRID. Having observed that a single rate is oftentimes split up into two neighbouring rates in the GRID to model off-GRID values we raised this to a tolerance of 3 fields. We calculated the scalar product for 100 stochastic simulations with identical parameters to give the fraction of inferred results with matching spectrum, where a matching spectrum must have a scalar product larger than 0.5. To visualize the effective amplitudes in Figure 2e, we locally integrated over three adjacent lines in the spectrum.

### Cell culture and preparation

Cells were cultured and prepared as described in ^25^. Cells with stable integration of Halo-CDX2 under doxycycline-induced expression control (kind gift from the lab of David Suter, EFPL, Lausanne, Switzerland) were seeded one day before the measurement on a Delta-T glass bottom dish (Bioptechs, Pennsylvania, USA). Expression of Halo-CDX2 was induced by adding 10ng/ml doxycyclin to the medium four hours before imaging. Cells were stained with SiR-dye (kindly provided by Kai Johnson, EFPL, Lausanne, Switzerland) shortly before imaging according to the Halo-tag protocol (Promega).

### Live cell single molecule imaging and tracking

Single molecule fluorescence imaging was performed as described previously ^41^. In brief, light of a 638 nm laser (IBEAM-SMART-640-S, 150 mW, Toptica, Gräfelfing, Germany) was used to set up a highly inclined illumination pattern on a conventional fluorescence microscope (TiE, Nikon, Tokyo, Japan) using a high-NA objective (100x, NA 1.45,Nikon, Tokyo, Japan). Emission light had to pass a multiband emission filter (F72-866, AHF, Tübingen, Germany) and was subsequently detected by an EMCCD camera (iXon Ultra DU 897U, Andor, Belfast, UK). For time-lapse imaging, dark-times were controlled by an AOTF (AOTFnC-400.650-TN, AA Optoelectronics, Orsay, France).

Cells were prepared for imaging as detailed above and kept in DMEM medium at 37° during imaging for up to two hours of measurement time per dish. Single molecule spot detection and tracking was performed as described in ^41^. In brief, we detected potential single molecules based on their fluorescence intensity compared to background fluorescence. Localization was performed using a 2D Gaussian fit. Halo-TF molecules were identified as bound molecules if they did not leave a radius of 288 nm for 5 consecutive frames. Fluorescence survival time distributions were extracted from these tracking data.

## Acknowledgements

We thank Achim Popp for culturing and preparing cells. SiR dye was kindly provided by Kai Johnsson (Max Planck Institute for Medical Research, Heidelberg). The work was funded by the European Research Council (ERC) under the European Union’s Horizon 2020 Research and Innovation Program (No. 637987 ChromArch to J.C.M.G.), the German Research Foundation (CRC 1279 Project B05 and GE 2631/1–1 to J.C.M.G.) and the German Academic Scholarship Foundation (to M.R.). The authors thank the Ulm University Center for Translational Imaging MoMAN for its support.

## Contributions

M.R., J.H. and J.C.M.G. designed the study. J.H. developed GRID and performed simulations. M.R. performed and analyzed experiments. T.K. contributed to image analysis. J.H. quantified TF diffusion on DNA. J.C.M.G. supervised the study. J.C.M.G., J.H. and M.R. wrote the manuscript.

## Data availability

Data supporting the findings of this manuscript will be available from the corresponding author after publication upon reasonable request. All raw single particle tracking data will be freely available in Matlab file format after publication.

## Code availability

The GRID software is freely available. A MatLab version of GRID and GRID simulation packages are available at https://gitlab.com/GebhardtLab/grid.

